# Bifurcation and sensitivity analysis reveal key drivers of multistability in a model of macrophage polarization

**DOI:** 10.1101/2020.07.06.188722

**Authors:** Anna-Simone Josefine Frank, Kamila Larripa, Hwayeon Ryu, Ryan Snodgrass, Susanna Röblitz

**Author notes:** Email addresses (Anna-Simone Josefine Frank), (Kamila Larripa), (Hwayeon Ryu), (Ryan Snodgrass), (Susanna Röblitz).

## Abstract

In this paper, we present and analyze a mathematical model for polarization of a single macrophage which, despite its simplicity, exhibits complex dynamics in terms of multistability. In particular, we demonstrate that an asymmetry in the regulatory mechanisms and parameter values is important for observing multiple phenotypes. Bifurcation and sensitivity analyses show that external signaling cues are necessary for macrophage commitment and emergence to a phenotype, but that the intrinsic macrophage metabolism is equally important. Based on our numerical results, we formulate hypotheses that could be further investigated by *in vitro* experiments to deepen our understanding of macrophage polarization.

**2010 MSC:** 93A30, 34C23, 49Q12, 92C42, 37N25

## 1. Introduction

Macrophages are highly versatile immune cells which, among other roles, eliminate pathogens and damaged cells through phagocytosis. They play a critical role in innate immunity and help to initiate the adaptive immune response through antigen presentation and cytokine signaling. Due to their diverse functions and plasticity, macrophages are able to exhibit markedly different phenotypes, depending on the external signals they receive, e.g., microbial products, damaged cells, or cytokines. The continuum of macrophage activation and the diverse spectrum of pro- and anti-inflammatory phenotypes result in nuanced immune regulations [31].

A conceptual framework has been developed for the description of macrophage activation with two polar extremes being the most widely studied and best understood. On one end of the phenotype spectrum, M1-like macrophages are classically activated by the cytokine interferon *γ* (IFN*γ*) or by an endotoxin directly [30]. Once activated, M1-like macrophages release cytokines that inhibit the proliferation of nearby cells (including cancer cells) and initiate inflammation and an immune response.

At the other extreme, M2-like macrophages are induced by the interleukins (IL)-4 and −13, cytokines secreted by activated Th2 cells [16]. They tend to dampen inflammation and promote tissue remodeling and tumor progression, for example through pro-angiogenic properties [4], immunosuppression (e.g., IL-10 expression) [21], remodeling of the extracellular matrix, or promotion of metastasis [24].

Mixed phenotypes also exist, which share some (but not all) significant features with the M1- or M2-like phenotypes [1]. The existence of mixed phenotypes has been particularly demonstrated in the tumor microenvironment [43].

Macrophage polarization is mediated in part, through the canonical Janus- or TYK2-kinases (JAK)-Signaling signal transducers and activators of transcription (STAT) signaling pathway. Activation of STATs is primarily driven by ligand-stimulated cytokine receptors whereby STATs become phosphorylated at a critical tyrosine residue leading to their release from the receptor complex where they then cross the nuclear membrane and reach chromatin. There they bind specific cognate DNA elements and participate in complex gene regulation processes.

Therefore, the phenotype expressed by a macrophage is identified through the specific STAT activation. M1 polarization is associated with STAT1 activity, whereas M2 polarization is associated with STAT6 activity [29].

The M1 and M2 polarization process is dynamic and can be reversed under certain conditions. Individual macrophages can change their phenotype in response to local signaling cues [46, 22, 51]. This can be especially pronounced in the tumor microenvironment and manifests in tumor associated macrophages, which can demonstrate both pro-tumoral and anti-tumoral activities [36].

Therefore, a better understanding of the polarization process of macrophages has the potential to guide the development of targeted cancer therapy to redirect the polarization towards a tumor suppressing microenvironment [47, 51, 7].

Mathematical modeling is a useful tool to better understand macrophage polarization by validating or testing hypothesis, and making predictions about possible dynamics. To our knowledge, three previous studies based on ordinary differential equations (ODEs) have modeled macrophage polarization and plasticity [38, 32]. While the authors in [32] showed bistable dynamics of macrophage phenotypes when exposed to external signaling cues, the authors in [38] could show that after initial differentiation into M1 and M2, the M2 phenotype was ultimately dominating. Finally, the authors in [50] used a systems-level approach to present the complexity of signaling pathways and intracellular regulation which describe macrophage differentiation under IFN-*γ*, IL-4 signaling, and cell stress (hypoxia). With their model, the authors in [50] could replicate experimental results on macrophage phenotype markers and transcription factor regulations upon external perturbations, also for the tumor microenvironment.

All three models are built using generic formulations of self-stimulation and mutual inhibition, which are also common building blocks in immune cell differentiation models [5, 49]. Similar modelling approaches as for T-cell differentiation have been used for macrophages in e.g., [32, 38], as T-helper cells differentiate in a similar manner [27, 29].

Our goal is to use mathematical modeling to shed light on the polarization and regulatory signaling dynamics related to activation of macrophage phenotypes. We aim to build a simple model, which includes less parameters than the previous models by [32, 38, 50], but which shows similar complex dynamics.

We conduct bifurcation and stability analyses to study its dynamical diversity, and relate these dynamics to biological observations. In addition, Global Sensitivity Analysis (GSA) is employed to i) guide the model reduction and ii) to identify the most sensitive drivers of the system dynamics. Finally, sensitive parameters and input signals values are altered to study their effect on the dynamics.

For the rest of this paper, Section 2 describes our mathematical model in context of macrophage polarization and Section 3 contains the conduction of the numerical methods. In Section 4, our main results, consisting of bifurcation analysis (Sec. 4.1, GSA (Sec. 4.2 and perturbation analysis based on GSA results (Sec. 4.3, are presented. We conclude with the Discussion in Section 5. The Appendix section provides more details on numerical analysis and the applied methodology.

## 2. Mathematical Model

Our mathematical model is based on the interactions specific to the macrophage lineage commitment signaling network. For this purpose, we simplify the network of macrophage functions in the liver from [37], and consider only IFN*γ* (input signal *S*1) and IL-4 (input signal *S*2) as relevant cytokine signals. The levels of activated STAT1 (variable *x*_1_) and STAT6 (variable *x*_2_) are used in our model as proxies for the two macrophage activation states.

A schematic diagram of our model is given in Fig. 1. We model the dynamics of activated STATs with a pair of coupled nonlinear differential equations, described in equations in (1)–(2).

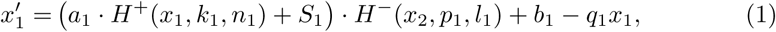

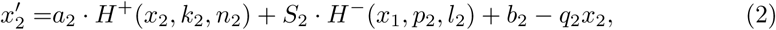

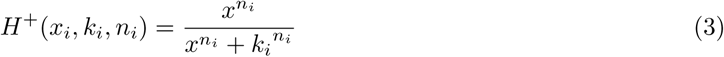

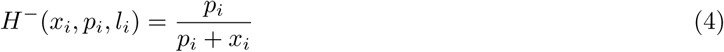

where 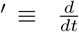. All parameters are assumed to be constant, positive and real numbers, except *n*_1,2_, which are integers.

**Figure 1:**
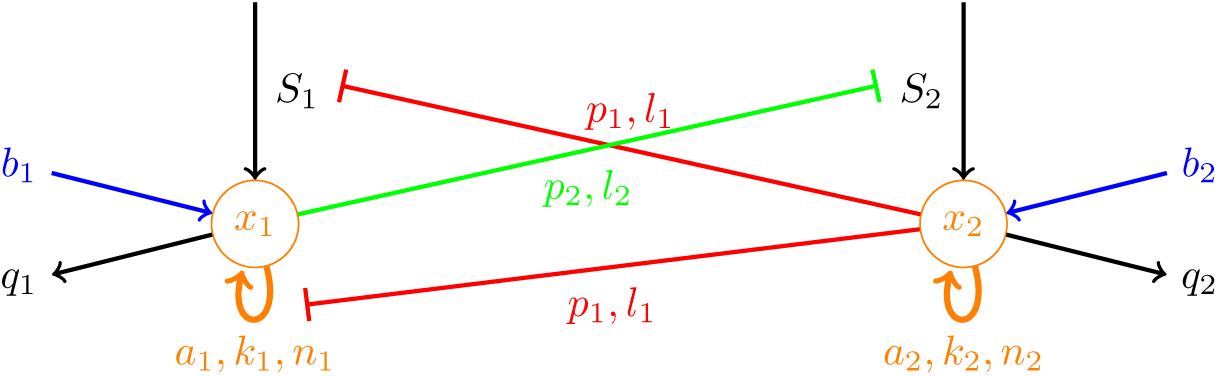
Schematic Diagram of Mathematical Model in equations (1)–(2). Self-stimulation of *x*_1_ and *x*_2_ are represented via the orange arrows, while processes of mutual-inhibition are shown by red and green inhibiting arrows. The incoming blue arrows depict *x*_1,2_ activation at basal rates (also in the absence of cytokine signaling), while the incoming black arrows represent the respective activation of *x*_1_ and *x*_2_ via cytokines (*S*_*i*_). Deactivation of *x*_1,2_ is illustrated by the outgoing black arrows.

The description of all model parameters is provided in Table 1.

**Table 1:**
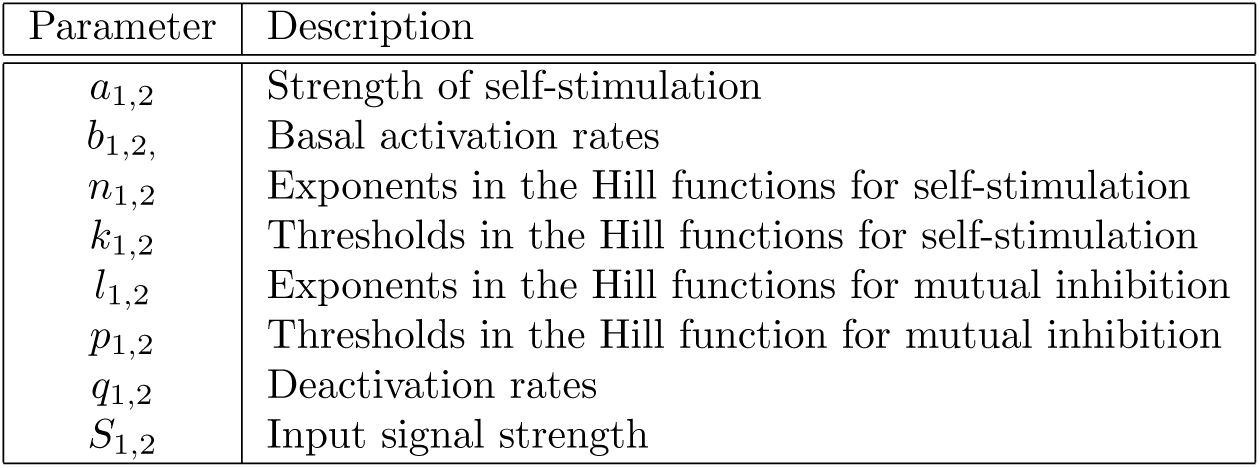
Model Parameters in equations (1)–(2)

| Parameter | Description |
| --- | --- |
| $a_{1,2}$ | Strength of self-stimulation |
| $b_{1,2}$ | Basal activation rates |
| $n_{1,2}$ | Exponents in the Hill functions for self-stimulation |
| $k_{1,2}$ | Thresholds in the Hill functions for self-stimulation |
| $l_{1,2}$ | Exponents in the Hill functions for mutual inhibition |
| $p_{1,2}$ | Thresholds in the Hill function for mutual inhibition |
| $q_{1,2}$ | Deactivation rates |
| $S_{1,2}$ | Input signal strength |

### 2.1. Model formulation

The equations in (1)–(2) were adapted from the T-cell model in [49]. The equation for *x*_2_ is based on the assumption that both type I and type II interferons inhibit IL-4-induced STAT6 activation in human monocytes in a SOCS-1-dependent manner [10], and therefore differs from the model formulation in [49]. This change results in an asymmetry in our equations in that STAT6 inhibits both the input signal and self-stimulation, but STAT1 affects only the input signal [44]. Furthermore, we reduced model complexity by fixing the Hill coefficient in equation (4) to 1. Also, the signal input function in [49] was simplified to a single parameter (*S*_1_, *S*_2_, respectively) for each phenotype in our model.

In our model equations, the parameters *α*_*i*_ represent the maximal activation rate of STAT due to self-stimulation. STATs are however also activated at low background levels (*b*_*i*_) in the absence of cytokine stimulation [9]. STATs are also inactivated by dephosphorylation, and we assume this rate is linear (terms *q*_*i*_*x*_*i*_ in the equations).

The fact that STAT1 and STAT6 are autocrine [48, 15], is captured by the stimulating Hill functions in the model equations (1)–(2). Finally, we assume respective activation of STAT1 and STAT6 via IFN*γ* (*S*_1_) and IL-4 (*S*_2_) [33].

We use stimulating (equation (3)) and inhibiting (equation (4)) Hill functions to describe STAT self-stimulation and mutual inhibition [42], respectively. The rationale behind the choice of these generic functions is that self-stimulation and inhibition are complex, non-linear processes, which consist of several individual steps. For example, in the process of self-stimulation, cytokines from the macrophage are secreted to stimulate helper T-cell differentiation [23]. Differentiated helper T-cells then secrete cytokines which in-turn stimulate the macrophage differentiation. However, detailed knowledge about these individual steps is unknown, which makes it difficult to derive mathematical equations for each step. In addition, we assume that the response in self-stimulation is sigmoidal, depending on the “dose” of input signals. Therefore, the Hill function is used and replaces the need to model the steps individually [42]. A similar argument was used for the inhibitory Hill function.

In the Hill function of equation (3), *k*_*i*_ represents the signaling level at which STAT stimulation is half-maximal and the Hill coefficient *n*_*i*_ governs the steep-ness of the Hill function in that as this value grows, the function becomes more switch-like. For the inhibitory Hill function, the parameters play a similar role.

## 3. Numerical Methods

In this section we provide the detailed description of numerical methods we employed.

### 3.1. Selection of model parameters

We explore parameter variations and analyze how the different parameter sets affect variability in the system states by using three parameter sets: the initial set **Θ**_0_, and two variation sets, **Θ**_1_ and **Θ**_2_. The parameters in the initial set **Θ**_0_ are adapted from [49], while the variation sets **Θ**_1_ and **Θ**_2_ are derived using nullclines. All three parameter cases are presented in Table 2.

**Table 2:**
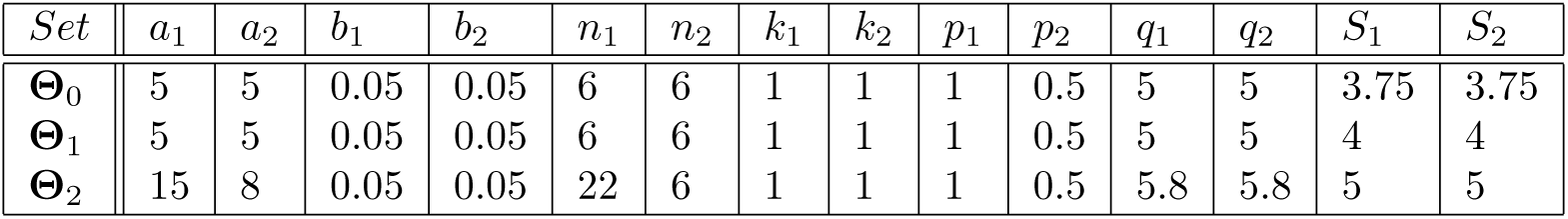
Parameter sets for numerical scenarios

| Set | $a_1$ | $a_2$ | $b_1$ | $b_2$ | $n_1$ | $n_2$ | $k_1$ | $k_2$ | $p_1$ | $p_2$ | $q_1$ | $q_2$ | $S_1$ | $S_2$ |
| --- | --- | --- | --- | --- | --- | --- | --- | --- | --- | --- | --- | --- | --- | --- |
| $\Theta_0$ | 5 | 5 | 0.05 | 0.05 | 6 | 6 | 1 | 1 | 1 | 0.5 | 5 | 5 | 3.75 | 3.75 |
| $\Theta_1$ | 5 | 5 | 0.05 | 0.05 | 6 | 6 | 1 | 1 | 1 | 0.5 | 5 | 5 | 4 | 4 |
| $\Theta_2$ | 15 | 8 | 0.05 | 0.05 | 22 | 6 | 1 | 1 | 1 | 0.5 | 5.8 | 5.8 | 5 | 5 |

### 3.2. Bifurcation and stability analysis

We expect our model, for all three case scenarios, **Θ**_0,1,2_, to exhibit at least bistable dynamics, similar to the original model. Thus, we first conduct bifurcation analysis to further investigate the impact of different parameter sets on model dynamics.

Bifurcation analysis aims to detect critical points of the bifurcation parameters, where the system dynamics change qualitatively in the long-term [17]. Given the biological importance of external signaling cues (INF-*γ* and IL-4) in the macrophage polarization process [46], we are primarily interested in determining how the system dynamics change based on varying input signals (i.e., *S*_1_ and *S*_2_). We therefore consider *S*_1_ and *S*_2_ as main bifurcation parameters, with the other parameters set to their values in Table 2. The bifurcation diagrams from equations (1)–(2) were obtained using the software package XPPAUT [12]. Details on numerical settings to draw bifurcation diagrams can be found in Appendix A.1.

We define states of STAT activation based on model-specific thresholds. An activation level is defined as *low*, if *S*_1,2_ ≤ 1.0, and as *high*, if *S*_1,2_ *>* 1.0. It is then the ratio of STAT1 to STAT6 activation, that characterizes a macrophage phenotype. The threshold levels are chosen to allow a consistent classification of phenotype cases in our model, although they only represent relative levels.

Stability analysis was performed by numerical simulations in Matlab.

### 3.3. Sensitivity analysis

We perform sensitivity analysis to identify parameter sets that have the greatest influence on the model outputs (e.g., STAT1 and STAT6 activation), and act as key drivers of macrophage polarization. Local sensitivity analysis quantifies changes in the model with respect to perturbation of a single parameter at-a-time, but is not recommended for non-linear models [52]. In contrast to local sensitivity, GSA methods explore the effects of changes in parameter values on model outcome by varying all parameters simultaneously. Although several global methods exist (e.g., extended Fourier Amplitude Sensitivity Test, Latin Hypercube Sampling and Partial Rank Correlation Coefficient [45, 28]), we employ Sobol’s method [39], a variance decomposition technique. We chose this method because it makes no assumptions about the relationship between model inputs and outputs in contrast to, for example, the Partial Rank Correlation Coefficient method, which requires monotonicity. Additionally, Sobol’s method considers interactions between parameters. A detailed description of Sobol’s method can be found in Appendix Appendix A.2.

We implemented Sobol’s sensitivity analysis using the SALib package [18]. We varied parameters 15% in each direction from their baseline values (i.e., parameter sets **Θ**_**0**,**1**,**2**_ in Table 2. We consider these scenarios separately. In all cases, we generated 300, 000 parameter set samples. The selected outcome of interest for the analysis is the ratio of STAT1 to STAT6 activation, which is responsible for macrophage polarization to specific phenotypes.

### 3.4. Perturbation in sensitive parameters

Based on results of the GSA, we explore the effect of perturbations in sensitive parameters on macrophage polarization dynamics. For illustrative purposes, we will only consider perturbations in the most sensitive parameter (*q*_2_) on case **Θ**_**0**_. Understanding the effect of dephosphorylation on system dynamics is especially important as deactivation rates change often in biological settings [19]. By changing *q*_2_ and keeping all other parameters fixed, we hope to better understand its individual effect on the relation between external input signals and activation of transcription factors.

## 4. Model Results

The results of the numerical simulations are presented in this section.

### 4.1. Bifurcation and stability analysis reveal multistable macrophage phenotypes

We observe bistability, tristability, and quadstability for different combinations of the *S*_1_ and *S*_2_ based on the three parameter cases **Θ**_0,1,2_, respectively.

#### 4.1.1. Bistable case

With the initial parameter set **Θ**_0_ we observe two stable fixed points, exhibiting bistable behavior. These steady states represent state variable ratios (*x*_1_*/x*_2_) with i) high/low and ii) low/low levels.

Though the detailed bifurcation diagrams are omitted here, we validate this bistable behavior by numerically solving equations (1)–(2) with the parameter set **Θ**_0_. The most interesting behavior observed is that *x*_1_ and *x*_2_ go through a switch before converging to their respective stable fixed points, as shown in Fig. 2(a). The solution trajectory of this switch behavior (in solid black) in the phase plane is provided in Fig. 2(b). Note that only two fixed points are present even though there seems to be another fixed point on the upper left part in the phase plane because of the proximity of the *x*_1_- and *x*_2_-nullclines. The bistable behavior is further confirmed by the basin of attraction shown in Fig. 2(c).

**Figure 2:**
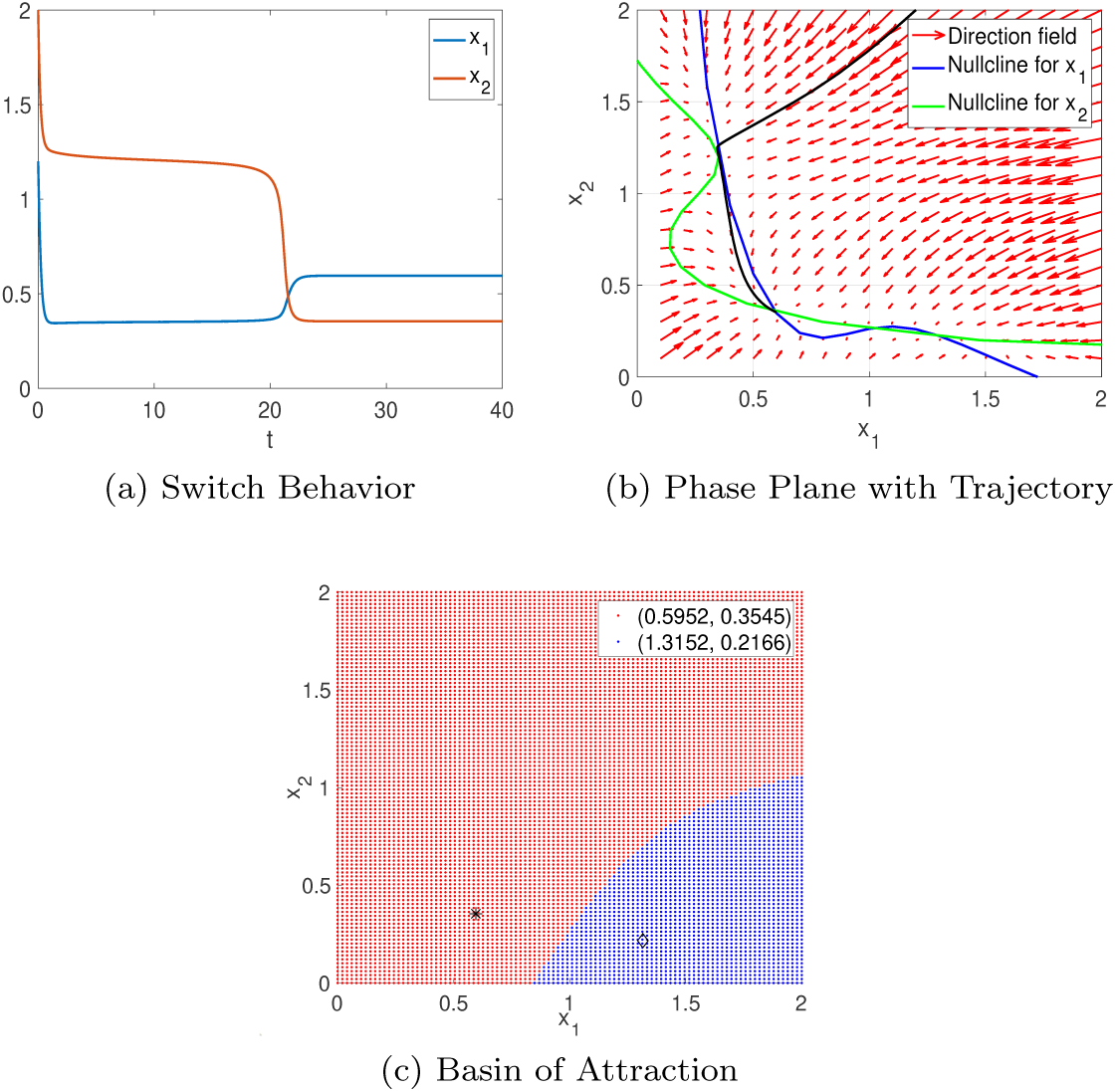
(a): Numerical solution that converges to low/low steady state after switch with initial condition (*x*_1_, *x*_2_) = (1.2, 2); (b): its corresponding trajectory (in solid black) in the phase plane; (c): the basin of attraction for both stable fixed points.

#### 4.1.2. Tristable case

With parameter set **Θ**_1_, three stable steady states of (*x*_1_*/x*_2_) are observed with i) low/high, ii) high/low, iii) low/low levels. The third steady state represents a situation where both STAT1 and STAT6 are present at similar levels.

Numerical solutions that converge to different stable fixed points are shown in Figs. 3(a)–(c). The respective solution trajectories in the phase plane are shown in Fig. 3(d).

**Figure 3:**
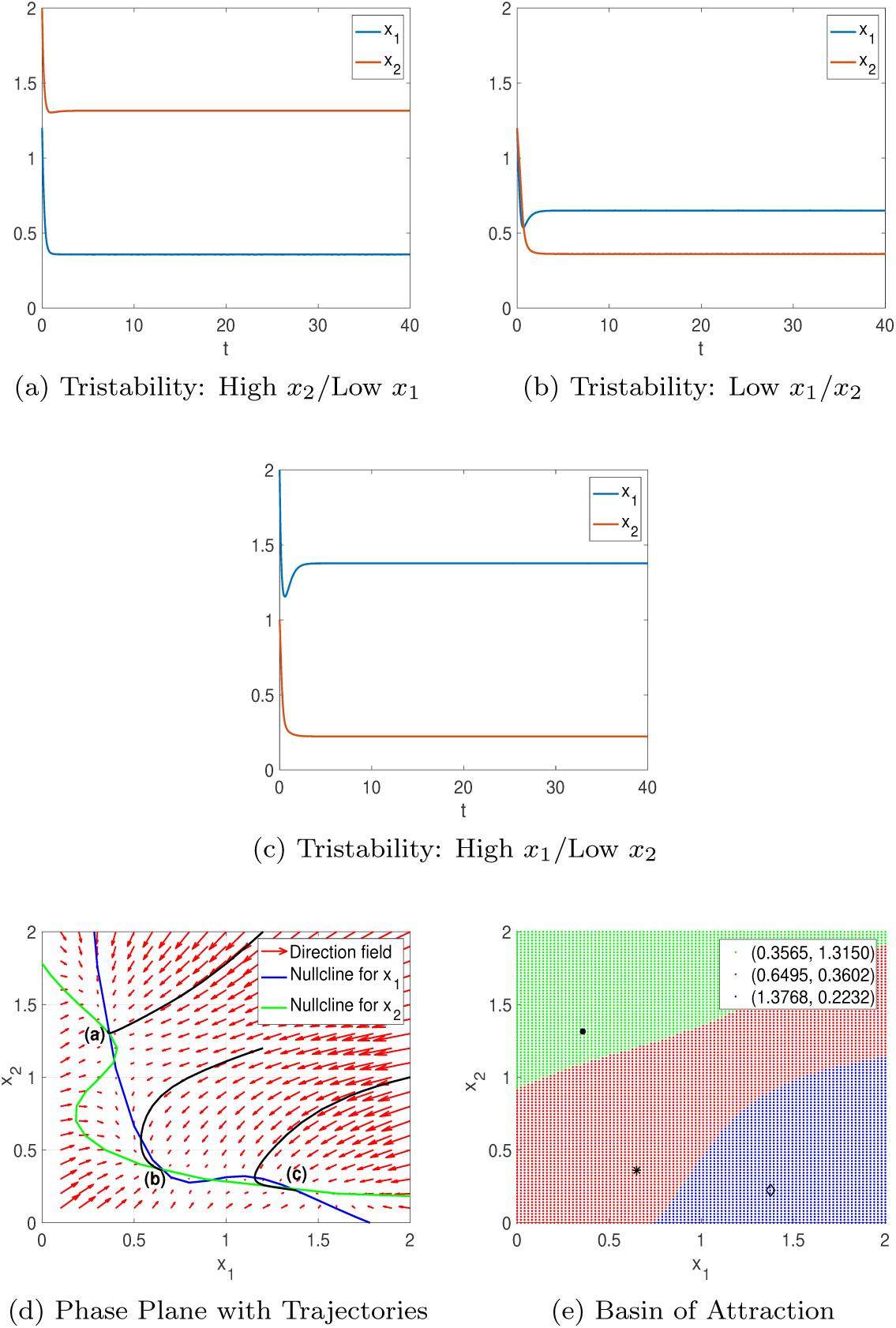
(a)–(c): Numerical solutions that converge to three different stable steady states with initial conditions (a) (*x*_1_, *x*_2_) = (1.2, 2), (b) (*x*_1_, *x*_2_) = (1.2, 1.2), and (c) (*x*_1_, *x*_2_) = (2, 1); (d): their corresponding trajectories (in solid black) in the phase plane; (e): the basin of attraction for each stable fixed point.

Because of the increased values *S*_1_ = *S*_2_ = 4 for this case, there are two additional intersections between the *x*_1_ and *x*_2_-nullclines compared to the bistable case, as can be seen in the phase plane of Fig. 3(d). This results in the addition of two fixed points, one of which is stable and the other is unstable. Thus, if we start with the same initial condition used in Fig. 2(a), the trajectory converges to the new stable fixed point with high *x*_2_/low *x*_1_, which was not observed in the bistable case. As further confirmed by the basin of attraction of Fig. 3(e), the other two stable fixed points remain as before.

It is the ratio of STAT1 (*x*_1_) to STAT6 (*x*_2_) activation levels that defines the polarization of a macrophage into the M1 or M2 phenotype [46, 32]. In our results, a high level of activated STAT1 in presence of low activated STAT6 levels defines the M1 phenotype [13], while low levels of activated STAT1 and high levels of activated STAT6 define the M2 phenotype. Low STAT1 and STAT6 activation levels represent a “hyporesponsive” phenotype that has not been described in the current literature. This phenotype might however have biological relevance (e.g., for cancer therapy), as an intermittent phenotype between M1 and M2. For example, recent studies by [3, 6, 25] have shown that tumors are initially characterized by M1 or an intermittent phenotype state, while advanced cancer is defined by M2 phenotype. It is therefore possible that this “hyporesponsive” phenotype describes another intermittent phenotype that appears during this transition.

#### 4.1.3. Quadstable case

Using the last parameter set **Θ**_2_, our model demonstrates quadstable behavior. The detailed bifurcation diagrams are provided in Fig. 4, where red solid lines represent stable fixed points, and black solid lines represent both unstable fixed points and saddle-nodes. Three of the stable fixed points, i.e., low/low, high/low and low/high, (in Figs. 4(a)–(d)) are qualitatively the same as those in the tristable case.

**Figure 4:**
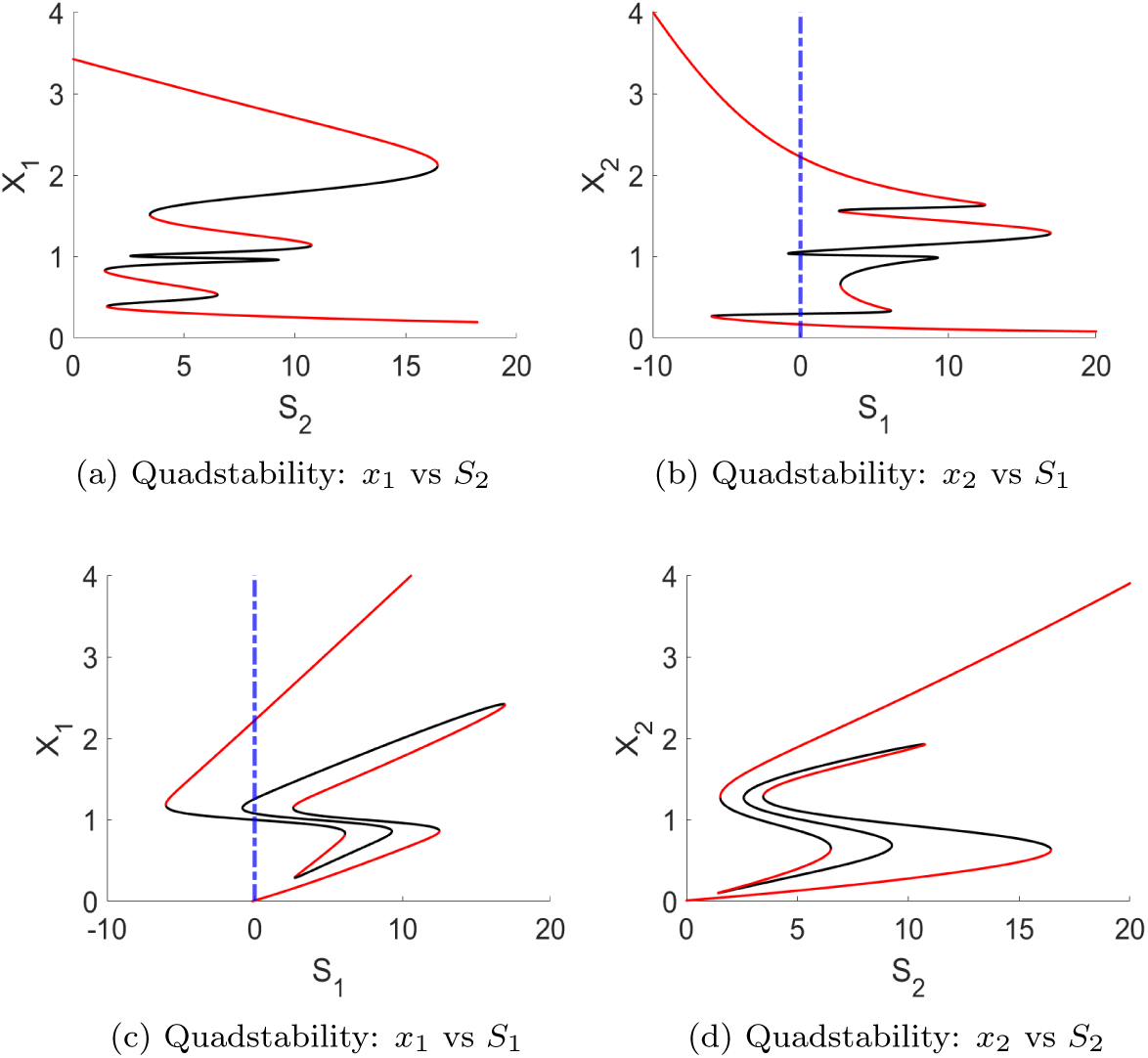
The bifurcation diagrams for varying input signals (*S*_1_ and *S*_2_) against the state variables *x*_1_ and *x*_2_ show quadstable dynamics (with the set **Θ**_2_). The red solid lines represent stable fixed points, while the black solid lines represent unstable fixed points and saddle-nodes. The blue dashed line represents the situation where *S*_1_ = 0.

The situation where both STAT1 and STAT6 have high activation status is, however, unique to the quadstable case. High activation of both STAT1 and STAT6 shows the existence of an intermittent phenotype [1], which bears characteristics of both the M1 and M2 types. Several of such intermittent states have been identified, for example, M2a, M2b, M2c and M2d [34]. The intermittent phenotype can also represent a transformation state, in which M1 branches to M2, and vice versa [8].

To understand how a varying input signal changes the activation of STAT1 and STAT6, we illustrate, based on Figs. 4(b)–(c), how one should read the bifurcation diagram: Figs. 4(b)–(c) have to be read simultaneously, starting from *S*_1_ = 0 and then increasing the *S*_1_ value while following the bifurcation trend. Note that while *S*_1_ is varied, all other parameters values are kept unchanged. By varying *S*_1_ from 0 to around 12, *x*_1_ is on the lowest stable branch while *x*_2_ is on the highest stable branch. Increasing *S*_1_ input signal beyond 12, *x*_1_ and *x*_2_ will follow the bifurcation trend up and down, respectively, to the next stable branch with *x*_1_ activation level between 1 and 2.2, and *x*_2_ activation level between 1.8 and 1.3. To reach the third stable branch, input signal *S*_1_ is decreased (to follow the bifurcation trend) until *x*_1_ and *x*_2_ jump from the second red branch to the third branch. The third branches spans values between 0.3 and 1 for *x*_1_, and values between 0.3 and 0.7 for *x*_2_. When on the third branch, *S*_1_ input signal will be increased again, at an input signal of around 7, both *x*_1_ and *x*_2_ will jump onto the respectively highest and lowest branch. Figs. 4(a)–(d) can be read similarly.

In Figs. 4(b)–(c), we observe furthermore that for high, *S*_1_ *>* 18 levels, the state variables *x*_1_ and *x*_2_ are committed to highest and lowest activation levels, respectively.

It is interesting that in the case of quadstability, the system is committed to the high/low state (see Figs. 4(b)–(c)) for high S1 values, while this could not be observed for bistable or tristable situations. Biologically, an irreversible switch into the M1 phenotype means that the macrophage will no longer be able to change its phenotype when exposed to changing input signals. This suggests that for high self-stimulation in the presence of high INF*γ* and low IL-4 signals, the system can commit to M1 phenotype and stay reversible for the M2 phenotype. In parameter set **Θ**_**2**_, STAT1 has higher self-stimulation than STAT6, i.e., *a*_1_ *> a*_2_ and *n*_1_ *> n*_2_. This might be a crucial driver for the commitment in the quadstable case, and the emergence of the intermittent phenotype.

Numerical solutions that converge to different stable points are shown in Figs. 5(a)–(d). Their respective solution trajectories are presented in Fig. 5(e). The basin of attraction of Fig. 5(f) shows the total of four stable fixed points, which indicates the quadstable dynamics.

**Figure 5:**
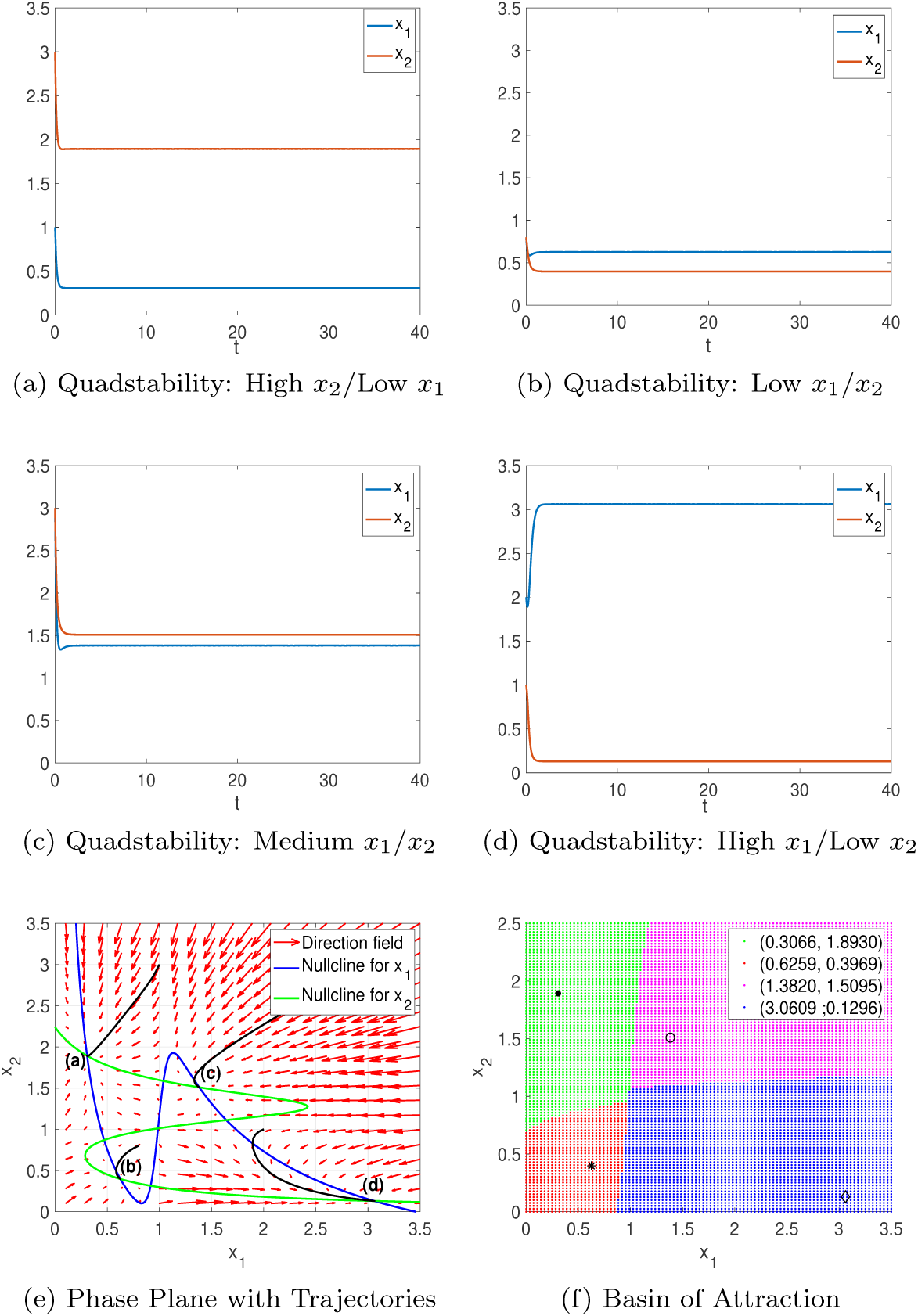
(a)–(d): Numerical solutions that converges to four different stable steady states with initial conditions (a) (*x*_1_, *x*_2_) = (1, 3), (b) (*x*_1_, *x*_2_) = (0.8, 0.8), (c) (*x*_1_, *x*_2_) = (3, 3), and (d) (*x*_1_, *x*_2_ = (2, 1); (e): their respective solution trajectories (in solid black) in the phase plane; (f): the basin of attraction for quadstable dynamics.

### 4.2. Identification of key drivers of macrophage dynamics through global sensitivity analysis

The most sensitive parameters for the bistable case using total sensitivity as a metric are, in descending order, *q*_2_, *q*_1_, *k*_2_, *S*_1_, *k*_1_, *a*_2_, *S*_2_ (see Fig. 6(a)). The four most sensitive parameters for bistable and tristable cases, shown in Figs. 6(a)–(b), respectively, agree and the next three most sensitive for each case are common (*k*_1_, *a*_2_, *S*_2_) but reordered. Fig. 6(c) shows that the most sensitive parameters in the quadstable case are consistent with results from the previous two cases.

**Figure 6:**
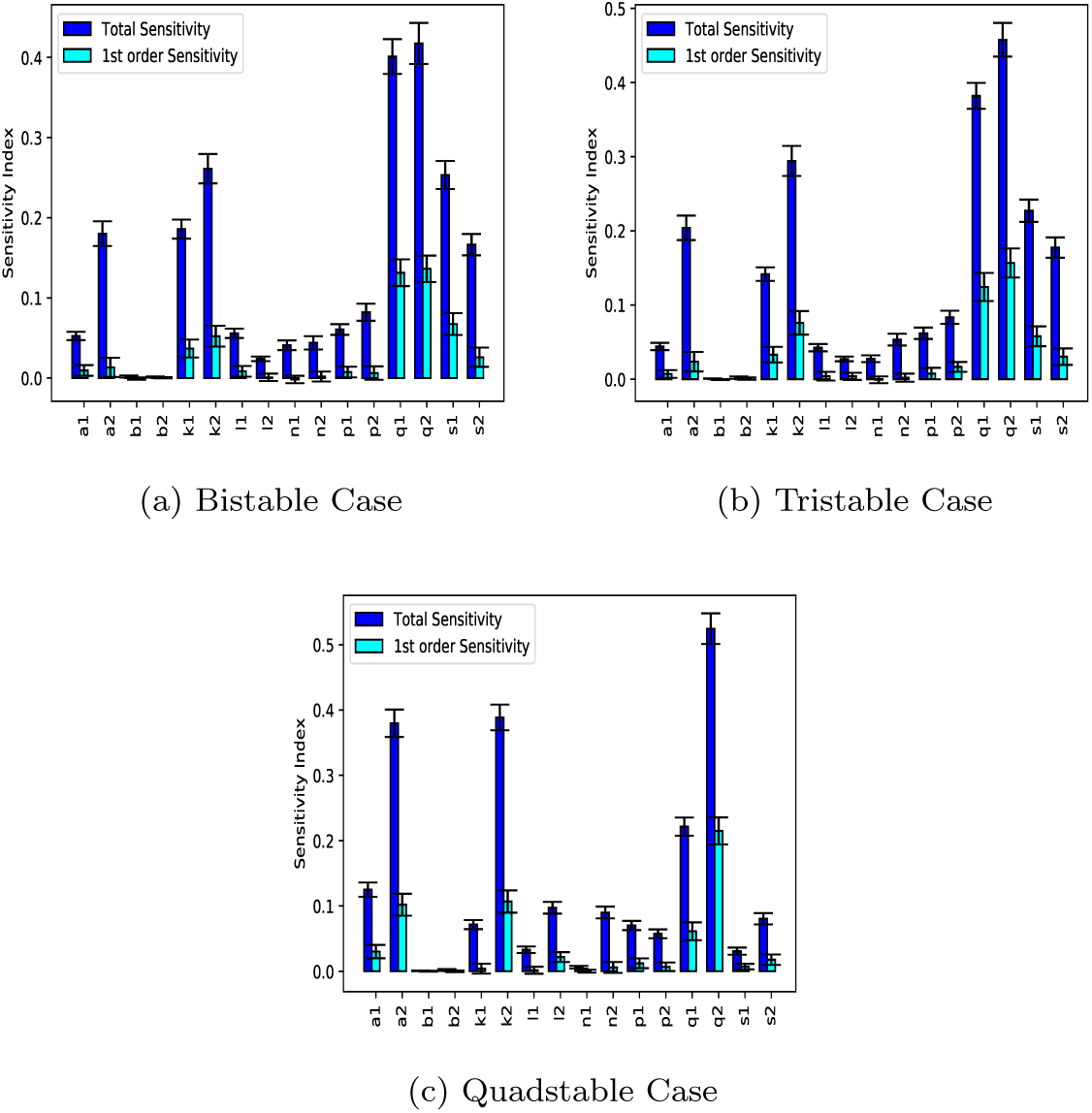
Sobol Sensitivity Indices where outcome of interest is the ratio of STAT1 activation to STAT6 activation at steady state. This used baseline parameter values which give (a) bistable, (b) tristable and (c) quadstable dynamics. In all instances, the parameter *q*_2_ has the highest total sensitivity index. The cases of bistability and tristability have the same most sensitive seven parameters *q*_2_, *q*_1_, *k*_2_, *S*_1_, *k*_1_ *a*_2_, *S*_2_ with only the ordering of the last three altered. For the quadstable case, *q*_2_ is also the most sensitive, with *k*_2_ and *a*_2_ moving up in the ordering compared to the previous two cases.

In terms of the pathways, this indicates that deactivation rates of both STAT1 and STAT6 (*q*_1_ and *q*_2_, respectively) are highly sensitive, as well as the input signal for M1 polarization, INF*γ* (*S*_1_). Parameters *k*_2_ and *k*_1_ are also sensitive, and both relate to the response of the Hill functions for self-stimulation. These parameters govern the concentration at which the switch takes place. In all cases, *k*_2_ is more sensitive than *k*_1_. Parameters *S*_2_ and *a*_2_ are the signaling input for M2 polarization (IL-4) and the maximum rate at which STAT6 stimulates its own activation via a regulative feedback mechanism.

### 4.3. Effect of perturbation in sensitive parameters

Figure 7 illustrates that by perturbing *q*_2_ the response of transcription factors to input signals changes. The change in response seems to occur with respect to the strength of the input signal, as well as according to stability. For example, in Figs. 7(b)–(c), lower *q*_2_ values seem to increase the number of stable states, and to increase the external stimuli needed to evoke a fate change. This example indicates that deactivation rates can contribute to the robustness of the dynamical system to variations in external stimuli. In particular, it illustrates that deactivation of STAT1 and STAT6 plays an essential role in macrophage polarization, as deactivation rates indirectly affect inhibition of external input signals on the opposite state variable, while self-stimulation affects its own state variable.

**Figure 7:**
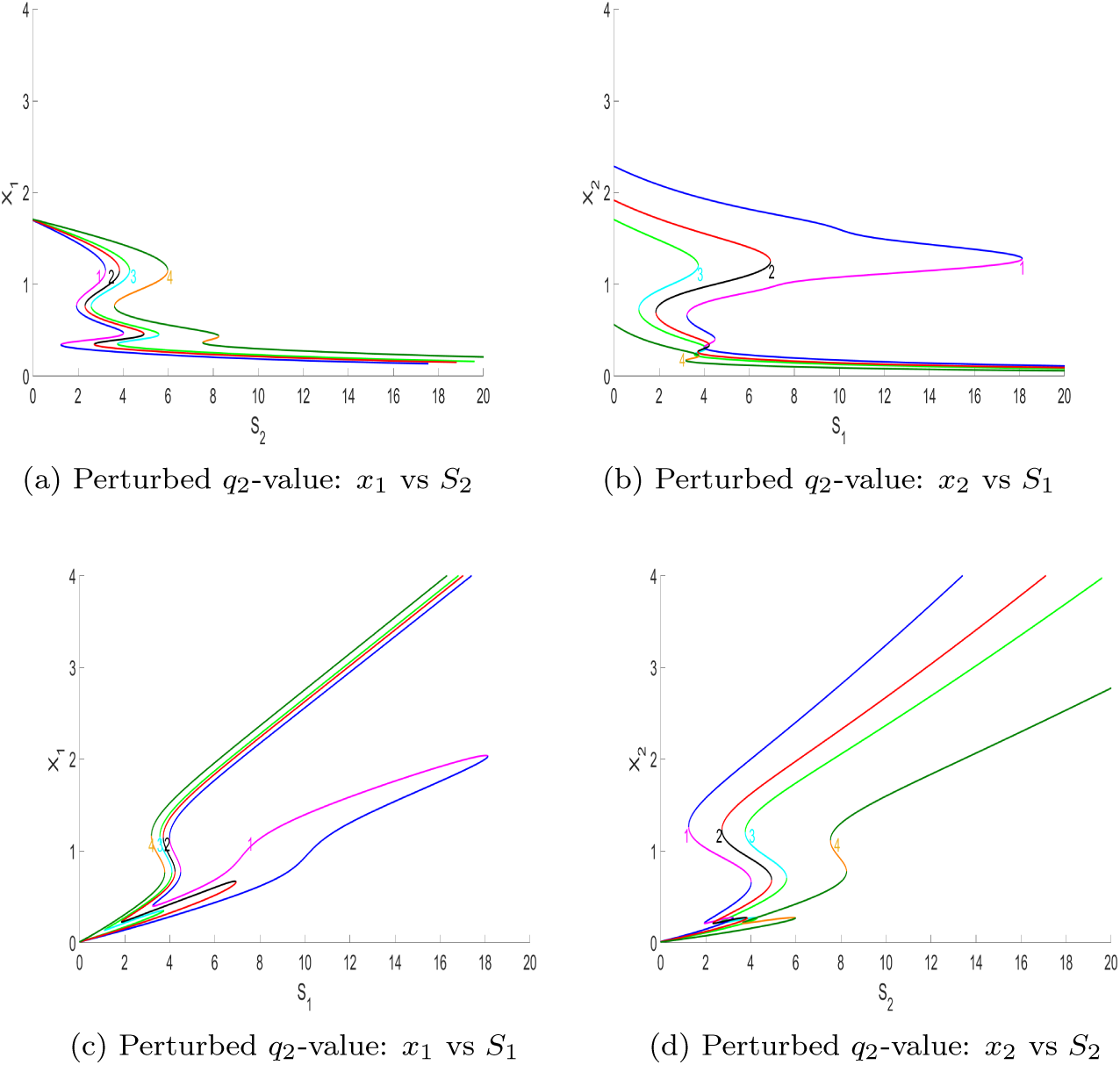
Case **Θ**_0_ for varying *q*_2_-values (*q*_2_ = 3.8–label 1, *q*_2_ = 4.5–label 2, *q*_2_ = 6.9–label 4) with respect to baseline *q*_2_-value (*q*_2_ = 5–label 3). The colors, magenta, black, turkeys and orange represent unstable branches, while blue, red, light green and dark green represent stable ones.

## 5. Discussion

In this work, we develop and explore a novel mathematical model for the dynamics of macrophage polarization and identify key parameters of the multi-stable dynamics. We validate that macrophage polarization is not strictly bipolar, but can consist of multiple phenotypes. Our macrophage polarization model is the first to show that a two-dimensional and simple model can predict bistable, tristable and quadstable phenotypes.

We could validate known phenotypes (i.e., M1 and M2) and have uncovered unknown, intermittent ones (i.e., low/low and high/high) with a mixed phenotype expression. From a biological perspective, the intermittent phenotypes might be more likely in *in vivo* settings than the extreme M1 and M2 cases, which are studied in cell cultures. Although, we cannot rule out that there exist more than four different phenotypes for our system, our findings are supported by those in [26], where the authors identified maximal four stable states given a similar model formulation. To our knowledge, only one previous study by [32], which studied a more complex model and applied also two- and three-dimensional bifurcation analyses, could identify a broader spectrum of known (e.g., M2a and M2b) and unknown macrophage phenotypes. Our identified unknown phenotypes can however not be compared directly to those in [32], because the authors classified STAT activation into high, medium and low levels, while we only made a distinction between high and low. In addition, such classification states are model dependent.

Sensitivity analysis of our model revealed the high impact of the deactivation rates, *q*_2_ and *q*_1_, on the ratio of STAT1 to STAT6 activation at steady state, used as a proxy for M1 and M2 phenotype, respectively. Parameters *k*_2_ and *α*_2_ were also identified as sensitive because both of these parameters are related to the self-stimulation of STAT6 activation. Our most sensitive model parameters are similar to those identified in [41, 50]. These sensitive parameters agree with results of our bifurcation analysis, where parameters of self-stimulation and deactivation seemed to have a profound impact on the dynamics. For example, in the quadstable case, parameters of self-stimulation might explain the observed system commitment and the emergence of an additional phenotype, while results of varying deactivation rates changed the response to external signaling cues, as can be seen in Fig. 7. The consistency in identifying sensitive parameters from bifurcation and sensitivity analyses is however expected, because a properly designed analysis should reveal bifurcation parameters to be sensitive [28].

In summary, bifurcation and sensitivity analyses showed that external signaling cues are necessary for macrophage commitment and emergence to a phenotype, but that the intrinsic macrophage metabolism (represented by self-stimulative factors and deactivation) is equally important [14, 2]. It should be noted that the intrinsic metabolic pathways, which enabled fate commitment in the quadstable situation, are masked by the generic nature (i.e., Hill function) of our model. Intrinsic metabolic pathways in macrophages are in general variable [14], and one can distinguish, for example, between glucose, lipid, iron or amino acid metabolic pathways [2].

Our results support the expectation from the model diagram (Fig. 1 that the system’s outcome also depends crucially on the self-stimulation of *x*_2_. Because the equations are not symmetric (i.e., in the second equation the stimulatory and inhibitory Hill functions are additive, not multiplicative as in the first equation), the parameters associated with STAT6 have a stronger impact on the model outcome. This observation is also reflected in the asymmetric values of *a*_1_, *n*_1_ and *a*_2_, *n*_2_ in **Θ**_**2**_. The asymmetry illustrates that lower values of *a*_2_, *n*_2_ have the same effect on systems dynamics as higher values of *a*_1_, *n*_1_. The parameters in **Θ**_**0**,**1**_ however are symmetric, because they were adapted from the mathematical model in [49], which has a symmetric model structure. The need for an asymmetry in self-stimulation dynamics of STAT1 and STAT6 might be explained by the experimental finding that the signaling pathway induced by IFN dominates over the signaling pathway induced by IL-4, according to the authors in [35]. This explanation is furthermore in accordance with our finding of an irreversible switch to the M1 phenotype for high concentrations of INF-*γ*.

Furthermore, we illustrated how STAT deactivation impacts macrophage polarization by influencing the robustness to external stimuli. The authors in [40] pointed out that the effects of deactivation are, however, not well understood for macrophages. Finally the knowledge of sensitive parameters for macrophage polarization might guide the conduction of future *in vitro* experiments and thus deepen our understanding of macrophage polarization. We therefore suggest that the following hypotheses, which resulted from our analyses, to be tested experimentally:

H1: The response-time and sensitivity of STATs to cytokine signaling levels can be altered by changing deactivation rates.

H2: Once macrophages are committed to a phenotype, further stimulation via cytokines leaves them unchanged.

H3: Intrinsic metabolic characteristics, which correspond to aspects of self-stimulation and deactivation, determine the range and variability of observable macrophage phenotypes.

H4: There exist intermittent phenotypes with equal STAT activation levels (i.e., defined by STAT phosphorylation) in *in vitro* settings.

### 5.1. Model limitations and future work

A clear advantage of our model is its simplicity and its ability to exhibits complex dynamics in terms of multistability. The price of simplicity is, however, the elimination of several key biological signaling components. One example is NF-*κ*B, a protein complex which interacts with type 1 interferons, among other signals [11]. Future work could inspect a more refined signaling network, based on our model formulation.

Another limitation of this work is that our model considers a single macrophage whereas in reality there are entire populations of macrophages which influence each other. However, understanding how a single macrophage reacts to its microenvironment is a first step to understanding population level behavior.

Mathematical models are needed to address macrophage polarization on population level and to consider input signals beyond IFN-*γ* and IL-4, while incorporating knowledge of dynamics of a single macrophage.

Finally, the absence of empirical data in this manuscript could be considered a limitation, especially as there exist data on the activation status of STATs. The primary focus of this manuscript has been to understand the qualitative characteristics of the proposed model. Hence extending the analysis to include empirical data is beyond this scope.

Our model represents also a solid first step towards analyzing stochastic gene expression in macrophages. In future work, we will make use of the chemical master equation and analyze how switching probabilities between different phenotypes change with variations in extrinsic and intrinsic noise levels.

## Acknowledgments

The authors thank the American Institute of Mathematics for providing support through its Structured Quartet Research Ensembles program.

## Data statement

This manuscript uses no biological or experimental data. All Figs. can be reproduced using the mathematical model, the parameters and numerical specifications presented in this manuscript.

## Appendix

### A. Appendix

#### Appendix A.1. Numerical details for bifurcation diagrams

Table A1 shows the numerical details used to calculate the bifurcation diagrams.

**Table A1:**
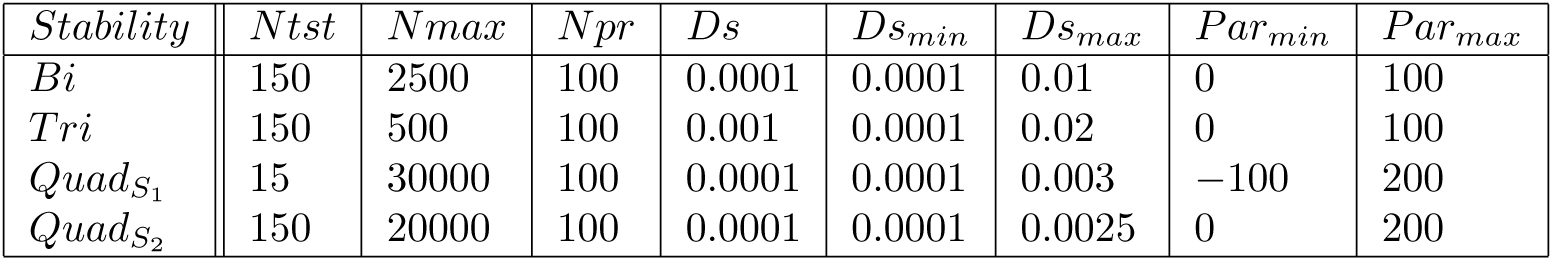
Numeric values for drawing bifurcation diagrams in XPPAUT. The column labels represent settings for numerical parameters in AUTO1.

| *Stability* | *Ntst* | *Nmax* | *Npr* | *Ds* | *Ds<sub>min</sub>* | *Ds<sub>max</sub>* | *Par<sub>min</sub>* | *Par<sub>max</sub>* |
| --- | --- | --- | --- | --- | --- | --- | --- | --- |
| *Bi* | 150 | 2500 | 100 | 0.0001 | 0.0001 | 0.01 | 0 | 100 |
| *Tri* | 150 | 500 | 100 | 0.001 | 0.0001 | 0.02 | 0 | 100 |
| *Quad<sub>S<sub>1</sub></sub>* | 15 | 30000 | 100 | 0.0001 | 0.0001 | 0.003 | −100 | 200 |
| *Quad<sub>S<sub>2</sub></sub>* | 150 | 20000 | 100 | 0.0001 | 0.0001 | 0.0025 | 0 | 200 |

1 Details can also be found at

http://www.math.pitt.edu/~bard/bardware/tut/xppauto.html

Abbreviations: *Ntst*, number of mesh intervals for discretization of periodic orbits, *Nmax*, maximum number of steps taken along any branch, *Npr*, give complete info every Npr steps, *Ds*, initial step size for bifurcation calculation, *Ds*_*min*_, minimum step size, *Ds*_*max*_, maximum step size, *Par*_*min*_, left-hand limit of the diagram for principal parameter, *Par*_*max*_, right-hand limit of the diagram for the principal parameter

#### Appendix A.2. Sobol’s method

Model output *f* (*x*) is decomposed into the sums of variances [39]:

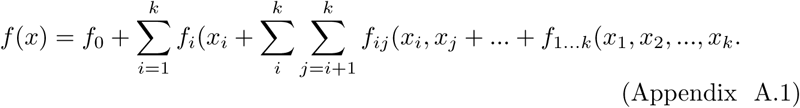

Here, *f*_*i*_ is the effect of varying *x*_*i*_ alone (first-order sensitivity), and *f*_*ij*_ is the effect of varying *x*_*i*_ and *x*_*j*_ simultaneously, additional to the effect of their individual variations, termed a second-order sensitivity. Higher order terms have analogous interpretations.

Assuming that *f* (*x*) is square integrable, the functional decomposition may be squared and integrated and the total variance D can be defined as

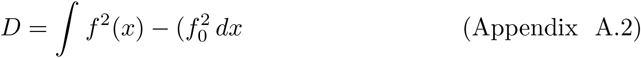

The partial variances from squaring and integrating the right hand side of Appendix A.1 are of the form

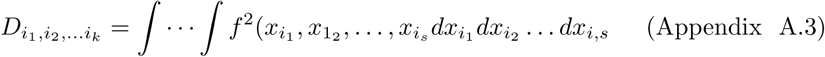

These integrals can then be approximated with Monte Carlo integration, and the Sobol sensitivity indices are calculated by the ratio of partial to total variance, representing the fraction of total variance which is attributed to individual model parameters or to combinations of parameters.

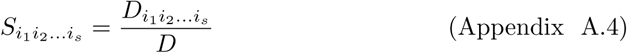

Furthermore, the total effect sensitivity index was proffered as an extension of the Sobol sensitivity index to quantify the overall effect of a parameter alone and in combination with any other parameters on model output [20]. This is defined to be

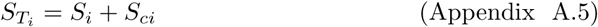

where *S*_*ci*_ is the set of sensitivity indices in which parameter *x*_*i*_ appears.

## References

[1] Biswas, S.K., Mantovani, A., 2010. Macrophage plasticity and interaction with lymphocyte subsets: cancer as a paradigm. Nat Immunol 11, 889–896. doi:10.1038/ni.1937.

[2] Biswas, S.K., Mantovani, A., 2012. Orchestration of metabolism by macrophages. Cell Metab 15, 432–437. doi:10.1016/j.cmet.2011.11.013.

[3] Bronte, V., Murray, P.J., 2015. Understanding local macrophage phenotypes in disease: modulating macrophage function to treat cancer. Nat Med 21, 117–119. doi:10.1038/nm.3794.

[4] Brown, J.M., Recht, L., Strober, S., 2017. The promise of targeting macrophages in cancer therapy. Clin Cancer Res 23, 3241–3250. doi:10.1158/1078-0432.CCR-16-3122.

[5] Callard, R.E., 2007. Decision-making by the immune response. Immunol Cell Biol 85, 300–305.

[6] Castiglione, F., Tieri, P., Palma, A., Jarrah, A.S., 2016. Statistical ensemble of gene regulatory networks of macrophage differentiation. BMC Bioinformatics 17, 506. doi:10.1186/s12859-016-1363-4.

[7] Cheng, H., Wang, Z., Fu, L., Xu, T., 2019. Macrophage polarization in the development and progression of ovarian cancers: an overview. Front Oncol 9, 421. doi:10.3389/fonc.2019.00421.

[8] Das, A., Sinha, M., Datta, S., Abas, M., Chaffee, S., Sen, C.K., Roy, S., 2015. Monocyte and macrophage plasticity in tissue repair and regeneration. Am J Pathol 185, 2596–2606. doi:10.1016/j.ajpath.2015.06.001.

[9] Dempoya, J., Matsumiya, T., Imaizumi, T., Hayakari, R., Xing, F., Yoshida, H., Okumura, K., Satoh, K., 2012. Double-stranded rna induces biphasic stat1 phosphorylation by both type i interferon (ifn)- dependent and type i ifn-independent pathways. J Virol 86, 12760–12769. doi:10.1128/JVI.01881-12.

[10] Dickensheets, H.L., Venkataraman, C., Schindler, U., Donnelly, R.P., 1999. Interferons inhibit activation of stat6 by interleukin 4 in human monocytes by inducing socs-1 gene expression. Proc Natl Acad Sci USA 96, 10800–10805. doi:10.1073/pnas.96.19.10800.

[11] Dorrington, M.G., Fraser, I.D., 2019. Nf-*κ*b signaling in macrophages: dynamics, crosstalk, and signal integration. Front Immunol 10, 705. doi:10.3389/fimmu.2019.00705.

[12] Ermentrout, B., 2001. XPPAUT 5.0-the differential equations tool. http://www.math.pitt.edu/~{}bard/xpp/xpp.html (accessed November, 2019).

[13] Fraternale, A., Brundu, S., Magnani, M., 2015. Polarization and repolarization of macrophages. J Clin Cell Immunol 6, 2. doi:10.4172/2155-9899.1000319.

[14] Geeraerts, X., Bolli, E., Fendt, S.M., Van Ginderachter, J.A., 2017. Macrophage metabolism as therapeutic target for cancer, atherosclerosis, and obesity. Front Immunol 8, 289. doi:10.3389/fimmu.2017.00289.

[15] Goenka, S., Kaplan, M.H., 2011. Transcriptional regulation by stat6. Immunol Res 50, 87–96. doi:10.1007/s12026-011-8205-2.

[16] Gordon, S., 2003. Alternative activation of macrophages. Nat Rev Immunol 3, 23–35. doi:10.1038/nri978.

[17] Gul, R., Bernhard, S., 2018. Sensitivity analysis: A useful tool for bifurcation analysis, in: Lopez-Ruiz, R. (Ed.), Complexity in Biological and Physical Systems: Bifurcations, Solitons and Fractals. Intech Open Limited, London, UK. chapter 4, p. 69. doi:10.5772/intechopen.72345.

[18] Herman, J., Usher, W., 2017. Salib: an open-source python library for sensitivity analysis. J Open Source Softw 2, 97. doi:10.21105/joss.00097.

[19] ten Hoeve, J., de Jesus Ibarra-Sanchez, M., Fu, Y., Zhu, W., Tremblay, M., David, M., Shuai, K., 2002. Identification of a nuclear stat1 protein tyrosine phosphatase. Mol Cell Biol 22, 5662–5668. doi:10.1128/mcb.22.16.5662-5668.2002.

[20] Homma, T., Saltelli, A., 1996. Importance measures in global sensitivity analysis of nonlinear models. Reliab Eng Syst Safe 52, 1–17. doi:10.1016/0951-8320(96)00002-6.

[21] Kuang, D.M., Zhao, Q., Peng, C., Xu, J., Zhang, J.P., Wu, C., Zheng, L., 2009. Activated monocytes in peritumoral stroma of hepatocellular carcinoma foster immune privilege and disease progression through pd-l1. J Exp Med 206, 1327–1337. doi:10.1084/jem.20082173.

[22] Lawrence, T., Natoli, G., 2011. Transcriptional regulation of macrophage polarization: enabling diversity with identity. Nat Rev Immunol 11, 750–761. doi:10.1038/nri3088.

[23] Lee, K.Y., 2019. M1 and m2 polarization of macrophages: a mini-review. Med Biol Sci Eng 2, 1–5. doi:10.30579/mbse.2019.2.1.1.

[24] Lin, E.Y., Nguyen, A.V., Russell, R.G., Pollard, J.W., 2001. Colonystimulating factor 1 promotes progression of mammary tumors to malignancy. J Exp Med 193, 727–740. doi:10.1084/jem.193.6.727.

[25] Linde, N., Gutschalk, C.M., Hoffmann, C., Yilmaz, D., Mueller, M.M., 2012. Integrating macrophages into organotypic co-cultures: a 3d in vitro model to study tumor-associated macrophages. PLoS One 7, e40058. doi:10.1371/journal.pone.0040058.

[26] Lu, M., Jolly, M.K., Gomoto, R., Huang, B., Onuchic, J., Ben-Jacob, E., 2013. Tristability in cancer-associated microrna-tf chimera toggle switch. J Phys Chem B 117, 13164–13174. doi:10.1021/jp403156m.

[27] Luckheeram, R.V., Zhou, R., Verma, A.D., Xia, B., 2012. Cd4+ t cells: differentiation and functions. Clin Dev Immunol 2012, 925135. doi:10.1155/2012/925135.

[28] Marino, S., Hogue, I.B., Ray, C.J., Kirschner, D.E., 2008. A methodology for performing global uncertainty and sensitivity analysis in systems biology. J Theor Biol 254, 178–196. doi:10.1016/j.jtbi.2008.04.011.

[29] Martinez, F.O., Gordon, S., 2014. The m1 and m2 paradigm of macrophage activation: time for reassessment. F1000Prime Rep 6, 13. doi:10.12703/P6-13.

[30] Medzhitov, R., 2008. Origin and physiological roles of inflammation. Nature 454, 428–435. doi:10.1038/nature07201.

[31] Mosser, D.M., Edwards, J.P., 2008. Exploring the full spectrum of macrophage activation. Nat Rev Immunol 8, 958–969. doi:10.1038/nri2448.

[32] Nickaeen, N., Ghaisari, J., Heiner, M., Moein, S., Gheisari, Y., 2019. Agent-based modeling and bifurcation analysis reveal mechanisms of macrophage polarization and phenotype pattern distribution. Sci Rep 9, 12764. doi:10.1038/s41598-019-48865-z.

[33] Ohmori, Y., Hamilton, T.A., 1997. Il-4-induced stat6 suppresses ifngamma-stimulated stat1-dependent transcription in mouse macrophages. J Immunol 159, 5474–5482.

[34] Palma, A., Jarrah, A.S., Tieri, P., Cesareni, G., Castiglione, F., 2018. Gene regulatory network modeling of macrophage differentiation corroborates the continuum hypothesis of polarization states. Front Physiol 9, 1659. doi:10.3389/fphys.2018.01659.

[35] Piccolo, V., Curina, A., Genua, M., Ghisletti, S., Simonatto, M., Sabò, A., Amati, B., Ostuni, R., Natoli, G., 2017. Opposing macrophage polarization programs show extensive epigenomic and transcriptional cross-talk. Nat Immunol 18, 530–540. doi:10.1038/ni.3710.

[36] Saccani, A., Schioppa, T., Porta, C., Biswas, S.K., Nebuloni, M., Vago, L., Bottazzi, B., Colombo, M.P., Mantovani, A., Sica, A., 2006. p50 nuclear factor-*κ*b overexpression in tumor-associated macrophages inhibits m1 inflammatory responses and antitumor resistance. Cancer research 66, 11432–11440. doi:10.1158/0008-5472.CAN-06-1867.

[37] Sica, A., Invernizzi, P., Mantovani, A., 2014. Macrophage plasticity and polarization in liver homeostasis and pathology. Hepatology 59, 2034–2042. doi:10.1002/hep.26754.

[38] Smith, T.D., Tse, M.J., Read, E.L., Liu, W.F., 2016. Regulation of macrophage polarization and plasticity by complex activation signals. Integr Biol (Camb) 8, 946–955. doi:10.1039/c6ib00105j.

[39] Sobol, I.M., 2001. Global sensitivity indices for nonlinear mathematical models and their monte carlo estimates. Math Comput Simulat 55, 271–280. doi:10.1016/S0378-4754(00)00270-6.

[40] Sridharan, R., Cameron, A.R., Kelly, D.J., Kearney, C.J., O’Brien, F.J., 2015. Biomaterial based modulation of macrophage polarization: a review and suggested design principles. Mater Today 18, 313–325. doi:10.1016/j.mattod.2015.01.019.

[41] Torres, M., Wang, J., Yannie, P.J., Ghosh, S., Segal, R.A., Reynolds, A.M., 2019. Identifying important parameters in the inflammatory process with a mathematical model of immune cell influx and macrophage polarization. PLoS Comput Biol 15, e1007172. doi:10.1371/journal.pcbi.1007172.

[42] Tyson, J.J., Novák, B., 2010. Functional motifs in biochemical reaction networks. Ann Rev Phys Chem 61, 219–240. doi:10.1146/annurev.physchem.012809.103457.

[43] Umemura, N., Saio, M., Suwa, T., Kitoh, Y., Bai, J., Nonaka, K., Ouyang, G.F., Okada, M., Balazs, M., Adany, R., et al., 2008. Tumorinfiltrating myeloid-derived suppressor cells are pleiotropic-inflamed monocytes/macrophages that bear m1-and m2-type characteristics. J Leukoc Biol 83, 1136–1144. doi:10.1189/jlb.0907611.

[44] Venkataraman, C., Leung, S., Salvekar, A., Mano, H., Schindler, U., 1999. Repression of il-4-induced gene expression by ifn-*γ* requires stat1 activation. J Immunol 162, 4053–4061.

[45] Wang, A., Solomatine, D.P., 2019. Practical experience of sensitivity analysis: Comparing six methods, on three hydrological models, with three performance criteria. Water 11, 1062. doi: https://doi.org/10.3390/w11051062.

[46] Wang, N., Liang, H., Zen, K., 2014. Molecular mechanisms that influence the macrophage m1–m2 polarization balance. Front Immunol 5, 614. doi:10.3389/fimmu.2014.00614.

[47] Williams, C.B., Yeh, E.S., Soloff, A.C., 2016. Tumor-associated macrophages: unwitting accomplices in breast cancer malignancy. NPJ Breast Cancer 2, 15025. doi:10.1038/npjbcancer.2015.25.

[48] Yarilina, A., Park-Min, K.H., Antoniv, T., Hu, X., Ivashkiv, L.B., 2008. Tnf activates an irf1-dependent autocrine loop leading to sustained expression of chemokines and stat1-dependent type i interferon–response genes. Nat Immunol 9, 378–387. doi:10.1038/ni1576.

[49] Yates, A., Callard, R., Stark, J., 2004. Combining cytokine signalling with t-bet and gata-3 regulation in th1 and th2 differentiation: a model for cellular decision-making. J Theor Biol 231, 181–196. doi:10.1016/j.jtbi.2004.06.013.

[50] Zhao, C., Mirando, A.C., Sové, R.J., Medeiros, T.X., Annex, B.H., Popel, A.S., 2019. A mechanistic integrative computational model of macrophage polarization: Implications in human pathophysiology. PLoS Comput Biol 15, e1007468. doi:10.1371/journal.pcbi.1007468.

[51] Zheng, X., Turkowski, K., Mora, J., Brüne, B., Seeger, W., Weigert, A., Savai, R., 2017. Redirecting tumor-associated macrophages to become tumoricidal effectors as a novel strategy for cancer therapy. Oncotarget 8, 48436–48452. doi:10.18632/oncotarget.17061.

[52] Zi, Z., 2011. Sensitivity analysis approaches applied to systems biology models. IET Sys Biol 5, 336–346. doi:10.1049/iet-syb.2011.0015.

